# Extraction-free iDNA metabarcoding of small dung beetles is an efficient method for surveying arboreal mammals in tropical forests

**DOI:** 10.64898/2026.01.16.699964

**Authors:** Torrey W. Rodgers, Arianna Basto, Raider Castro, Erin Marcela Rivera Groves, Charles C. Y. Xu, Elena Hernández, Karen E. Mock, Adrian Forsyth

**Affiliations:** Ecology Center and Department of Wildland Resources, Utah State University Logan, Utah, USA; New Venture Fund, Washington DC, USA; Faculty of Biological Sciences, Ricardo Palma University, Lima, Peru; Andes Amazon Fund, Washington DC, USA; Health Program, Wildlife Conservation Society, Bronx, New York, USA; Faculty of Veterinary Medicine, University of Calgary, Alberta, Canada

**Keywords:** Amazon Basin, biodiversity monitoring, direct PCR, Invertebrate-derived DNA, Madre de Dios, Peru, Primate surveys, Scarabaeidae

## Abstract

Surveying mammals in tropical rainforests, particularly those that inhabit the rainforest canopy, can be challenging. We tested a novel method for detecting rainforest mammals with metabarcoding of iDNA at Los Amigos Biological Station in the Peruvian Amazon. We used an extraction-free, direct PCR approach to metabarcode vertebrate DNA, whereby buffer from tubes that contained dung beetles for 24 hours was used directly in PCR, negating the need for labor-intensive dissection and DNA extraction. We used a mammal-specific marker targeting the mitochondrial 16S gene and a general metazoan marker targeting the mitochondrial COI gene. From just 105 dung beetles, mostly small beetles from the genus *Sylvicanthon,* we detected 42 vertebrate species from 16 orders and 26 families, including 33 mammal species and 10 primates. Of the mammals, more than half were arboreal species. The mammal-specific 16S marker detected more mammal species, but the general COI marker detected some mammals not detected by 16S plus eight avian and one fish species, and, in most cases, yielded sequences from the dung beetles themselves. Because small dung beetles are often abundant and easy to catch in tropical forests, this extraction-free direct PCR method provides a powerful and efficient tool for monitoring mammalian diversity. Our results demonstrate that this method works particularly well for primates and other arboreal mammals that are challenging to detect with traditional methods.

## Introduction

Environmental DNA (eDNA) metabarcoding has become a valuable technique for monitoring biodiversity (Bohmann et al. 2014, Ruppert et al. 2019). While early applications of eDNA targeted aquatic vertebrates, recent advancements have extended the use of eDNA methods to terrestrial species, and invertebrate-derived DNA (iDNA) methods in particular have emerged as a powerful approach for detecting vertebrates (Newton et al. 2025, Broadhurst et al. 2025). Invertebrates such as carrion flies (Calvignac-Spencer et al. 2013, Rodgers et al. 2017), leeches (Ji et al., 2022) and dung beetles (Drinkwater et al., 2021; Saranholi et al., 2024), can act as eDNA samplers by collecting vertebrate DNA through interactions with animal carcasses, feces, or blood. Sequencing trace amounts of DNA derived from these interactions allows for indirect detection of vertebrates without visual observation or physical capture. Such non-invasive methods are particularly useful in complex environments like tropical rainforests, where high species richness, elusive animal behavior, dense vegetation, and diverse canopy-dwelling fauna may reduce the effectiveness of conventional survey tools like camera traps (Rodgers et al., 2017).

While iDNA metabarcoding is effective for vertebrate sampling (Rodgers et al. 2017, Ji et al. 2022, Saranholi et al. 2024, Broadhurst et al. 2025), further methodological improvements are needed to enhance efficiency and decrease labor and cost. Among the steps in eDNA and iDNA metabarcoding workflows, DNA extraction remains one of the most labor-intensive. Previous dung beetle iDNA metabarcoding studies have performed dissection of dung beetle digestive tracts followed by a traditional DNA extraction of the gut contents (Drinkwater et al. 2021, Saranholi et al. 2024), a process which demands considerable labor. A method that negates the need for labor-intensive dissection and DNA extraction would reduce laboratory processing time by many hours and greatly reduct cost. Thus, we tested an extraction-free, direct polymerase chain reaction (PCR) approach for next-generation sequencing of mammal DNA derived from dung beetles whereby buffer added to tubes that contained live beetles for 24 hours was used directly in PCR. Although this approach has previously been used with carrion flies (Srivathsan et al. 2023, Rodgers et al. 2025), this study marks its first application to dung beetles. Here, we demonstrate that this technique is highly effective for detecting mammalian DNA, allowing for efficient surveys of mammal communities in tropical forests.

Extraction-free Direct PCR was conducted with a mammal specific marker targeting the mitochondrial 16S gene. In addition, we investigated the utility of using a general metazoan cytochrome c oxidase I (COI) marker to detect mammals, as well as other vertebrate species, while simultaneously identifying the COI barcode sequences of the dung beetles themselves with direct PCR. We also tested the efficacy of non-invasively swabbing the bodies of large dung beetles for vertebrate DNA before releasing them. These methodological advancements streamline dung beetle iDNA metabarcoding workflows, thus expanding their utility for rapid, cost-effective, and non-invasive monitoring of vertebrate diversity in challenging tropical rainforest environments, especially for difficult to detect canopy-living species.

## Methods

### Study site

Los Amigos Biological Station (LABS; also known as CICRA in previous literature), is located at 12°33’40.0" S, 70°05’46.4" W in the Amazon Basin at an elevational range from 225 to 296 meters above sea level in the Madre de Dios region of southeastern Peru (Figure 1). The sampling area was at the convergence of the Los Amigos and Madre de Dios rivers. This area includes a range of habitats including mature old growth forest on terra firme terraces of ancient, weathered soils of low pH, as well as both mature and successional forests populated by dense stands of *Guadua* bamboo and *Mauritia flexosa* palm swamps found on the more fertile alluvial floodplains that are annually renewed by flooding. The region has a tropical humid climate with a rainy season from November to February and a dry season from June to September. Annual rainfall ranges between 2700 and 3000 mm, and the mean temperature ranges from 21°C to 26 °C. The region supports an intact mammalian community typical of lowland rainforests in the western Amazon that is currently free from hunting.

**Figure 1.**
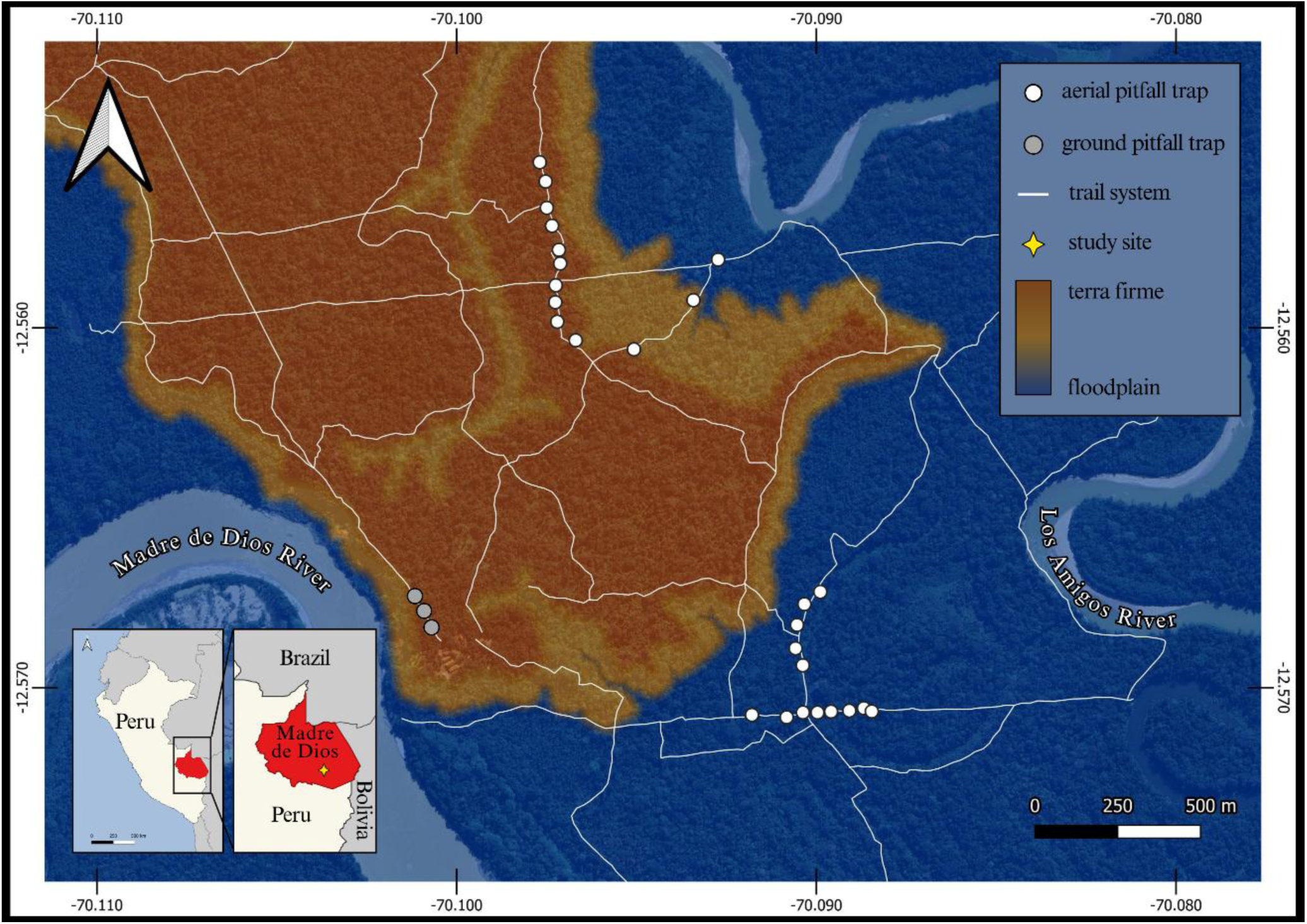
Map of the study site at Los Amigos Biological Station, Madre de Dios, Peru, showing locations where dung beetles were collected for iDNA metabarcoding to detect rainforest mammals.

### Dung beetle collection

We collected dung beetles using both aerial pitfall traps and ground pitfall traps baited with human feces. Beetles were collected with 10 aerial pitfall traps daily from Nov 5-11, 2023, and from 3 ground pitfall traps from Nov 9-10, 2023. We constructed aerial pitfall traps from plastic bottles (Fig 2), with bait wrapped in cloth hung by a string inside the top of the trap (Figure 2). A wire mesh was inserted into the plastic bottle to prevent beetles from accessing the bait directly. We placed aerial traps 50 m apart at a height of approximately two m by suspending them from trees with a rope. Ground pitfall traps were constructed of plastic bottles buried at ground level. We baited traps each morning at approximately 08:00, and we checked traps in the afternoon at approximately 14:00 and again the following morning at approximately 08:00. After three days of sampling, we moved aerial traps to a from tarra firme habitat to floodplain habitat. Ground pitfall traps remained at the same locations for both days. We placed beetles captured from each trap in bleach-sterilized resealable plastic bags to be transported back to the field laboratory.

**Figure 2.**
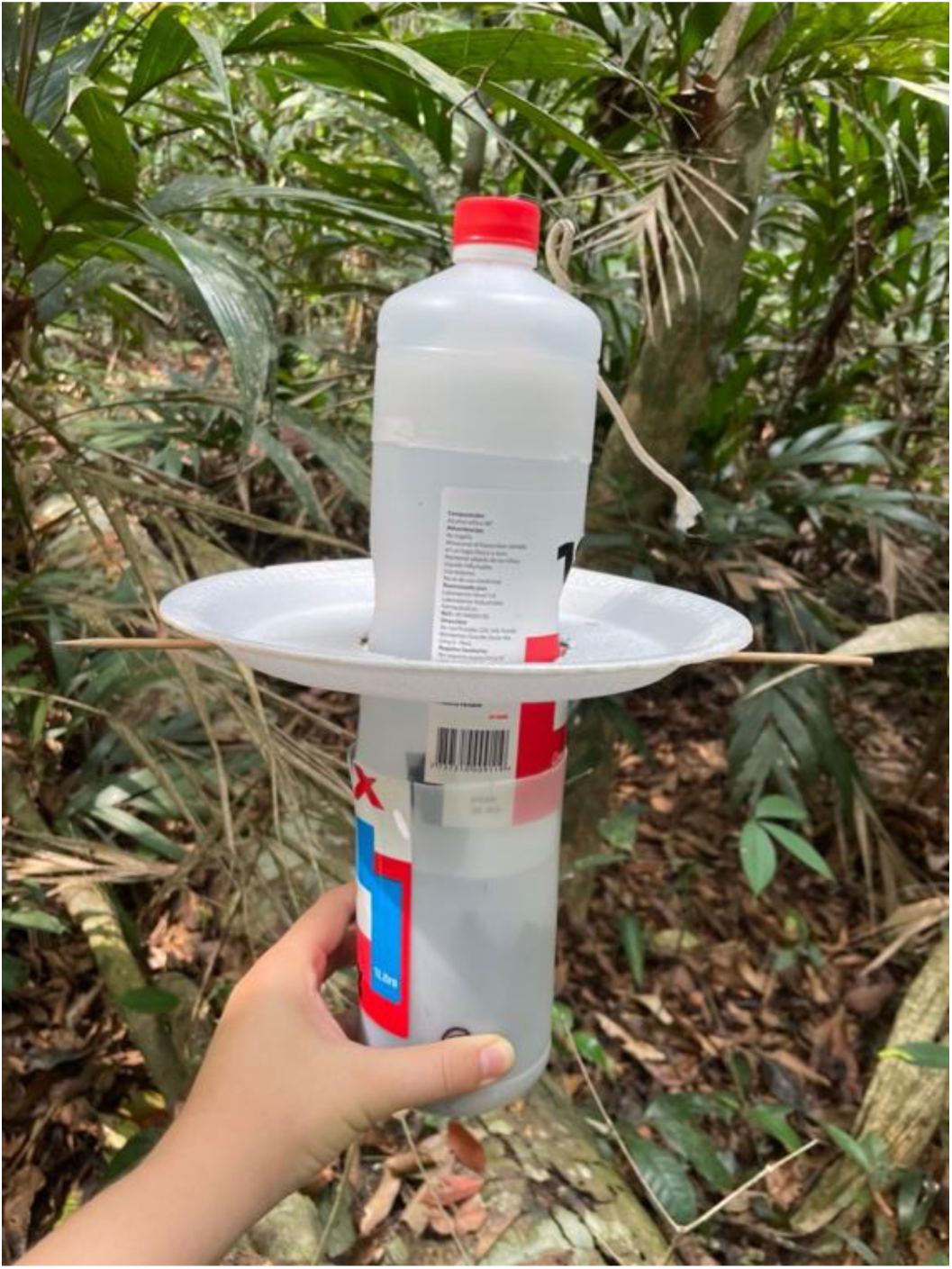
Arial pitfall trap used to collect dung beetles.

Once in the field laboratory, we placed individual live beetles in labelled two mL snap-cap microcentrifuge tubes with single-use latex gloves where they were left for approximately 24 hours (Figure 3A). We then removed beetles from tubes and placed them in whirl-pack bags with EtOH for morphological species identification. We also sampled DNA from seven beetles that were too large to fit into two mL tubes by swabbing their faces with a sterile cotton swab and then rotating the swab inside of a two mL tube to transfer DNA. Next, we placed 4-5 blue-indicating silica gel beads in each empty tube to dry the interior, and then we placed the tubes in a -20 °C freezer until transport to the molecular laboratory. Prior to transport, we removed and disposed of silica gel beads so that only empty, dry tubes were transported to the molecular ecology laboratory at Utah State University, where they were stored in a -70 °C freezer.

**Figure 3.**
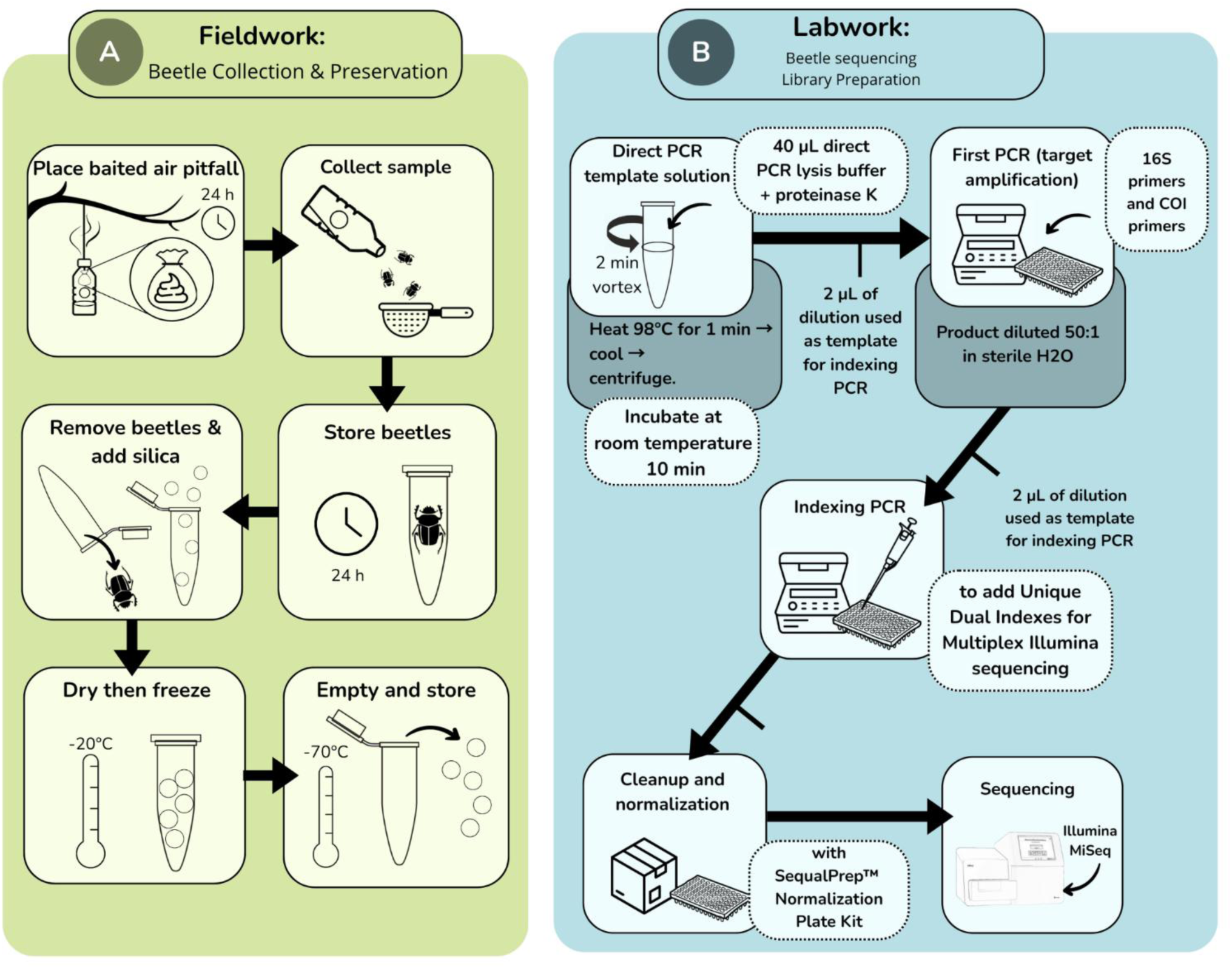
Workflow for A) dung beetle collection field work and B) laboratory work for sequencing of dung beetles for detection of rainforest mammals with iDNA from Los Amigos Biological Station, Madre de Dios, Peru.

**Figure 4.**
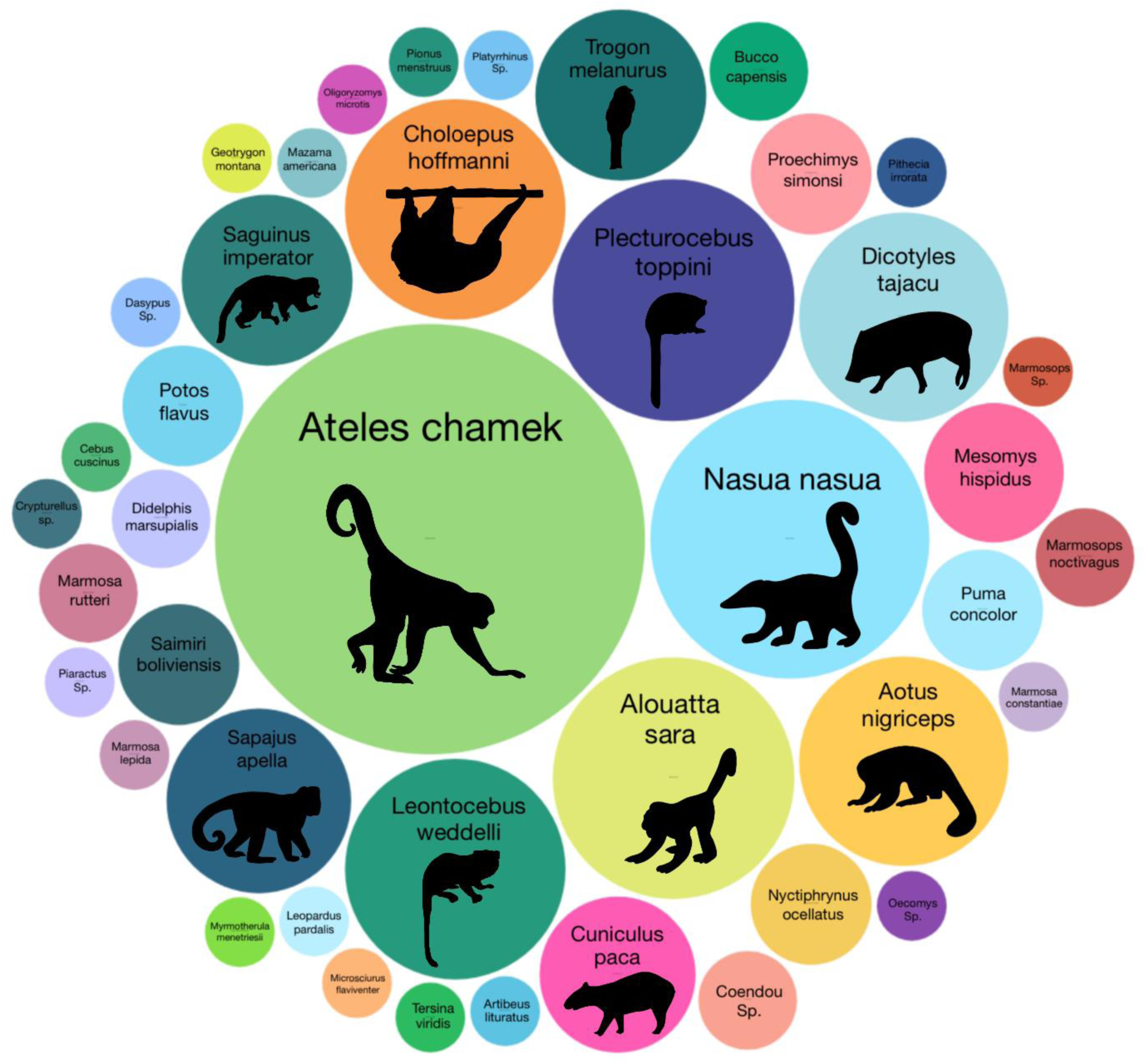
Visual representation of mammals detected by extraction-free metabarcoding of dung beetles from Los Amigos Biological Station, Madre de Dios, Peru. Size of circles represents the number of beetles containing DNA for each species (Table 3).

### Laboratory analysis

We used an extraction-free, direct PCR approach for DNA amplification (Figure 3B). We added 40 μL of direct PCR lysis buffer (supplied with Platinum™ Direct PCR Universal Master Mix, Invitrogen, Carlsbad, CA) with 1.2 μL of proteinase K to each microcentrifuge tube from beetle sampling. We vortexed tubes for two minutes, incubated them at room temperature for 10 minutes, and then incubated them on a heat block at 98 °C for one minute, before cooling to room temperature (as per manufacturers’ lysis protocol). We then centrifuged samples for one minute before adding the buffer template solution directly to PCR reactions.

We conducted PCR on all samples with two mitochondrial markers, one targeting the 16S rRNA gene, and another targeting the COI gene. PCR reactions for the 16S marker contained 10 μL Platinum™ Direct PCR Universal Master Mix (Invitrogen, Carlsbad, CA), 4 μL Platinum™ GC Enhancer, 0.2 μM each of forward and reverse primers (16Smam1-To and 16Smam2; Table 1) containing Illumina sequencing adapters, 1 μM (5X) human blocking primer (16Smam-blkhum; Table1), and 2 μL of direct PCR lysis buffer template solution from sample tubes for a final PCR reaction volume of 20 μL. PCR reactions for the COI marker contained 10 μL Platinum™ Direct PCR Universal Master Mix (Invitrogen, Carlsbad, CA), 4 μL Platinum™ GC Enhancer, 0.4 μM each of forward and reverse primers (mlCOIintF-exN and jgHCO2198; Table 1) containing Illumina sequencing adapters, and 2 μL of direct PCR lysis buffer template solution from sample tubes for a final PCR reaction volume of 20 μL. PCR conditions were: initial denaturation at 94 °C for 10 min, 40 cycles of 94 °C for 15 sec, 60 °C (16S) or 46 °C (COI) for 15 sec, and 68 °C for 20 sec, followed by a final elongation of 68 °C for 5 minutes. Each PCR run included a no-template control of sterile molecular grade water, a lysis buffer negative control, and a positive control containing DNA from bobcat *Lynx rufus*, a species that does not occur in South America. All negative and positive controls were indexed and sequenced.

**Table 1.**
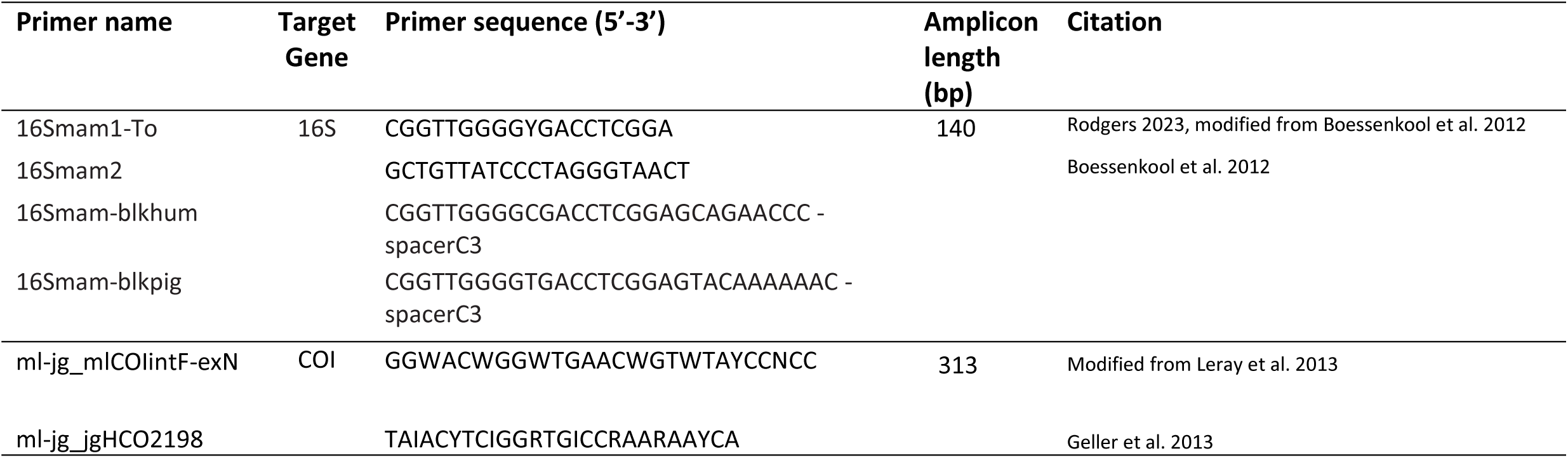
Primers used for metabarcoding of dung beetle iDNA samples.

We diluted PCR products 50:1 in sterile H_2_O for use as template in an indexing PCR to add unique dual indexes to each sample for multiplexing. Unique Dual Indexing (UDI) primers (Integrative DNA Technologies) contained the complimentary Illumina flow-cell adapter sequence and an 8 bp unique dual index on each end (forward and reverse) for unique sample identification from multiplexed samples. Indexing PCRs reactions included 10 μl of Amplitaq Gold DNA polymerase (Thermo Fisher Scientific), 0.5 uM of forward and reverse UDI indexing primers, and 2 μl of diluted PCR product from the first PCR in a total reaction volume of 20 μl. PCR cycling conditions for the indexing PCR were: initial denaturation at 95 °C for 10 min, 15 cycles of 95 °C for 15 sec, 50 °C for 30 sec, and 72 °C for 30 sec, followed by a final elongation of 72 °C for 7 minutes. Following the indexing PCR, we cleaned up and normalized PCR products using a SequalPrep™ normalization plate (Thermo Fisher Scientific) following the manufacturer’s protocol. We then pooled the resulting product and provided it for sequencing at the Center for Integrative Biosystems sequencing core at Utah State University. The 16S library was sequenced on an Illumina MiSeq with a 300 cycle v2 Micro reagent kit, and the COI library was sequenced on an Illumina NextSeq 2000 with a 600 cycle P1 reagent kit.

Bioinformatic processing of sequence data proceeded as follows. After demultiplexing, we removed primer sequences with CUTADAPT v. 1.18 (Martin 2011). Next, data were filtered, denoised, paired-ends were merged, and chimeras were removed with DADA2 (Callahan et al. 2016). Amplicon Sequence Variants (ASV) were merged into Operational Taxonomic Units (OTUs) at 97% similarity with VSEARCH (Rognes et al. 2016), all within the QIIME2 environment (Bolyen et al. 2019) version 2024.10.

For taxonomic assignment, we trained Naïve Bayes classifiers using scikit-learn (Pedregosa et al. 2011) within the QIIME2 environment. Classifiers were trained on lrRNA (16S) and COI reference databases comprising unique haplotypes from MIDORI2, v. GB266 (Leray et al. 2022). For 16S, the amplicon was first extracted from the database using our primer sequences with default parameters, after which the classifier was trained. For COI, because our reverse primer occurs in a conserved region that is commonly used for COI barcoding, many reference sequences within MIDORI2 lack this region because primer regions are often trimmed prior to submission, so we could not use the standard QIIME2 workflow for amplicon extraction. As an alternative, we used CUTADAPT to retain COI sequences containing only the forward primer with a match percentage of 0.8. We then removed all sequences shorter than 280 bp and truncated at 310 bp to create a uniform COI reference database for classifier training. In addition, for COI only, we used the python package BOLDigger3, v. 2.1.4 (Buchner and Leese 2020), a classifier designed specifically for taxonomic assignment of COI data using the Barcode of Life Database (BOLD; Ratnasingham & Hebert, 2007). In some cases, taxonomic assignment to the species level was made subjectively when our classifiers made assignments only to genus level or made a species level assignment to a species not known from Peru, but for which a member of the genus is known from Los Amigos based on local knowledge and a regional checklist (Payne et al. 2024). For example, we obtained a 16S OTU that our classifier assigned to the primate species *Pithecia pissinatti* with relatively low confidence (0.77) however this species is only known from Brazil. The congener *Pithecia irrorate* is known from our study area, however there is no 16S or COI reference sequence available for this rare species for assignment. Thus, we subjectively assigned this OTU to *Pithecia irrorate*.

## Results

### Dung beetle collection

We collected 105 individual dung beetles representing six genera and 13 species (Table 2). *Sylvicanthon proseni* was the most collected species (n=55), followed by *Sylvicanthon bridarolli* (n=18), with beetles from the genus *Sylvicanthon* representing 70% of all samples. Eighty-eight (84%) of beetles were collected from arial pitfall traps, and 17 (16%) were collected from ground pitfall traps.

**Table 2.** Dung beetle species collected for iDNA metabarcoding from Los Amigos Biological Station, Madre de Dios, Peru based on morphological identification. The total number collected, and the number and percentage of beetles from each species that yielded wild vertebrate DNA are listed.

| Species | Quantity | Yielded DNA |  |
| --- | --- | --- | --- |
|  |  | Number | Percent |
| *Sylvicanthon proseni* | 55 | 44 | 80% |
| *Sylvicanthon bridarolli* | 18 | 13 | 72% |
| *Canthon fulgidus* | 7 | 6 | 86% |
| *Onthophagus osculatii* | 7 | 5 | 71% |
| *Deltochilum amazonicum* | 5 | 3 | 60% |
| *Dichotomius batesi* | 4 | 4 | 100% |
| *Canthon quadrigutatus* | 2 | 2 | 100% |
| *Eurysternus caribaeus* | 2 | 2 | 100% |
| *Canthon luteicollis* | 1 | 1 | 100% |
| *Onthophagus osculati* | 1 | 1 | 100% |
| *Onthophagus transithmius* | 1 | 1 | 100% |
| *Eurysternus hamaticollis* | 1 | 1 | 100% |
| *Dichotomius conicollis* | 1 | 0 | 0% |
| Total | 105 | 83 | 79% |

### Laboratory analysis

Illumina MiSeq sequencing with the 16S marker resulted in 1,759,785 reads, of which we retained 654,250 after quality filtering, denoising, and chimera removal, with a median of 5,855 reads per sample. Illumina NextSeq sequencing with the COI marker resulted in 82,975,482 reads, of which we retained 15,401,825 after quality filtering, denoising, and chimera removal, with a median of 134,197 reads per sample.

We clustered reads from 16S into 59 unique OTUs, all from the phylum Chordata. Of these, 58 OTUs were assigned to class Mammalia, and one OTU was assigned to class Actinopteri with the Naïve Bayes classifier Scit-learn. After removal of contaminant OTUs identified as belonging to humans or domestic species, the *Lynx rufus* positive control, and OTUs not identified at family level, 34 unique OTUs from vertebrate species remained. We assigned these 34 OTUs to 26 unique taxa, 25 mammals and one fish. This included 10 primates, four carnivores (two felids and two procyonids), six rodents, two artiodactyls, two chiropterans, and one sloth (Table 3). Of these 26 unique taxa, 23 were assigned at the species level to a species known from Los Amigos, while the remaining three taxa were assigned only to genus.

**Table 3.**
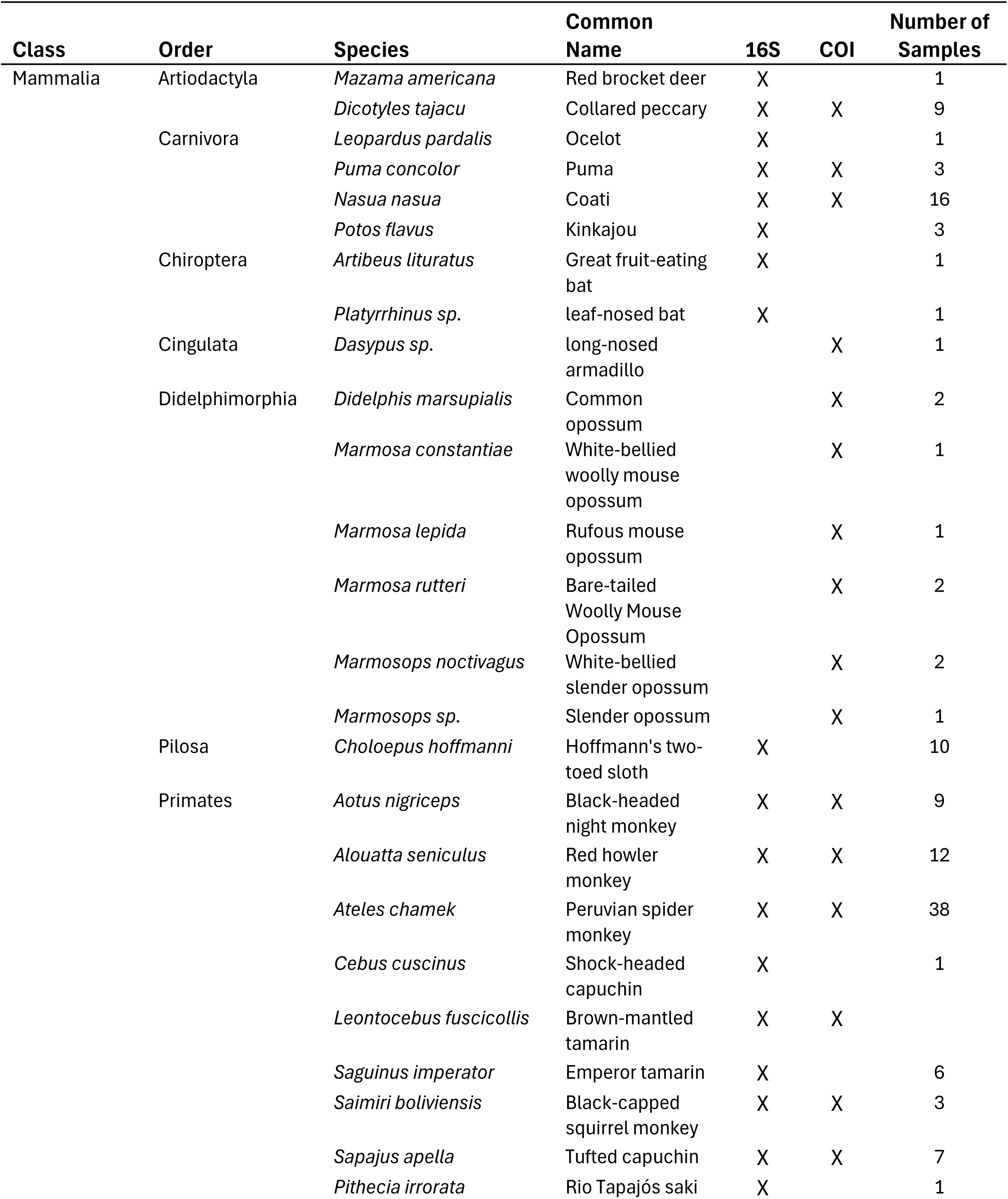
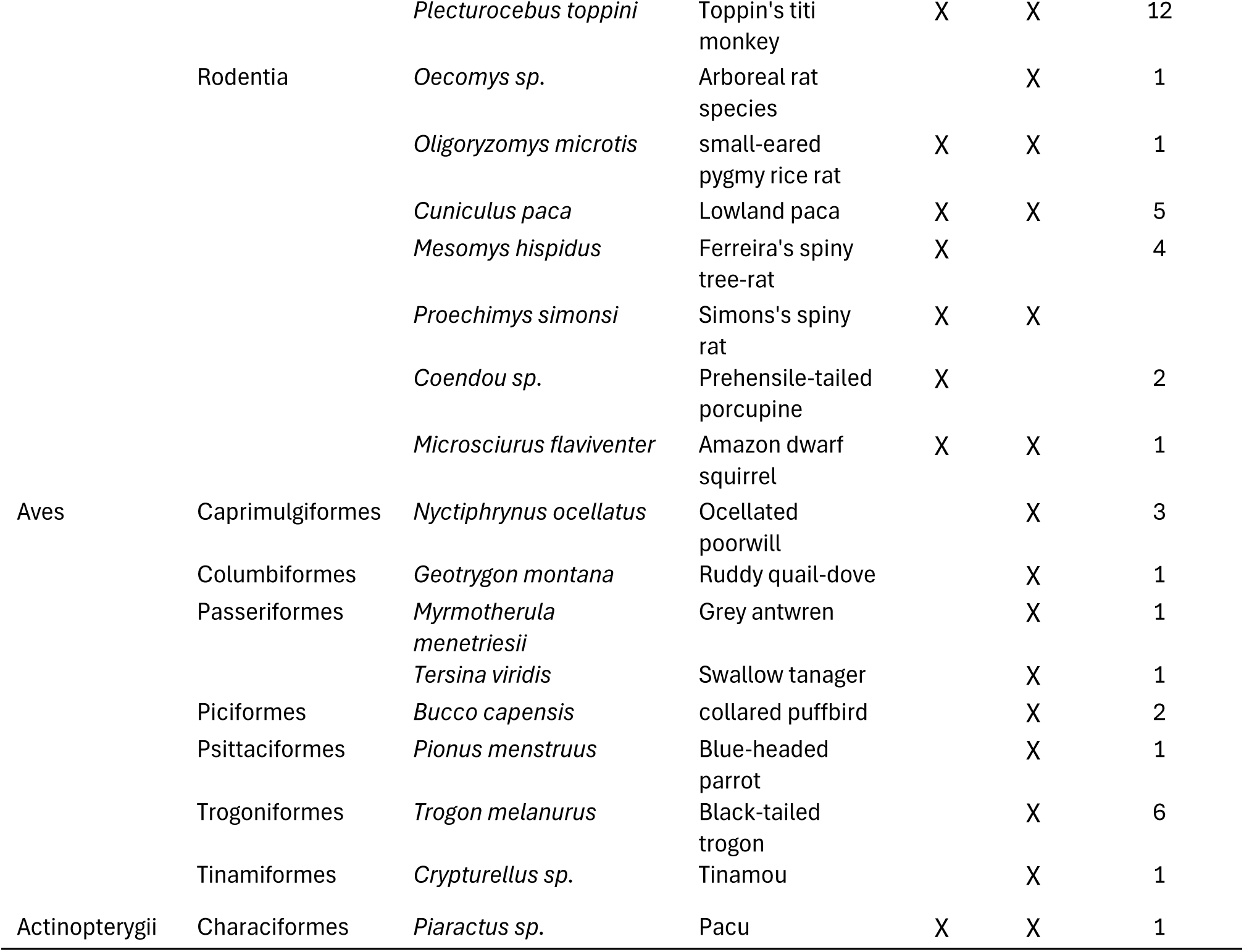
Vertebrate taxa detected from extraction-free iDNA metabarcoding of dung beetles from Los Amigos Biological Station, Madre de Dios, Peru with two different mitochondrial markers targeting the 16S rRNA gene and the cytochrome c oxidase I (COI) gene.

We clustered reads from COI into 48,030 unique OTUs. For the Naïve Bayes classifier Scit-learn, 89% of OTUs were assigned to phylum, with the remaining 11% only assigned to kingdom Eukaryota. Of those assigned to phylum, 29,346 OTUs (69%) were assigned to fungal phyla, with 12,835 (30%) assigned to animal phyla and the remaining 1% assigned to a variety of other phyla including amoebozoa, bacteria, algae and dinoflagellates. Among the animal phyla, 11,543 OTUs (90%) were assigned to Arthropoda, and 400 (3.3%) to Chordata, with the remaining 7% assigned to nematodes annelids and rotifers. After removal of OTUs identified as belonging to humans or domestic species and those not identified at family level, 111 mammalian OTUs, 19 avian OTUs and one fish OTU remained. These OTUs were assigned to 20 mammalian taxa, eight avian taxa, and one fish taxon (Table 3). Of the OTUs assigned to Arthropoda, 45 OTUs were assigned to family Scarabaeidae (dung beetles). The remaining arthropod OTUs were assigned to 17 orders and 70 families from the classes Arachnida, Collembola, and Insecta (Supplemental Table S1).

For Boldigger3, 5,074 OTUs (10.6%) were assigned at the phylum level, with the remaining 89% not assigned. Of those assigned to phylum, 2,278 OTUs (45%) were assigned to fungal phyla and 2,671 (53%) assigned to animal phyla with the remaining 2% assigned to assigned to a variety of other phyla including amoebozoa, bacteria, algae and dinoflagellates. Among the animal phyla, 1,873 OTUs (70%) were assigned to Arthropoda, and 666 (25%) to Chordata, with the remaining 5% assigned to nematodes annelids and rotifers. Of the OTUs assigned to Chordata, 641 were assigned to class Mammalia, 24 were assigned to class Aves, and one was assigned to class Actinopterygii.

After removal of OTUs identified as belonging to humans or domestic species and those not identified at family level, 89 mammalian OTUs, 7 avian OTUs and 1 fish OTU remained. These OTUs were assigned to 22 mammalian taxa, seven avian taxa, and one fish taxon (Table 3). Of the OTUs assigned to Arthropoda, 240 OTUs were assigned to the family Scarabaeidae (dung beetles). The remaining arthropod OTUs were assigned to 17 orders and 104 families from the classes Arachnida, Collembola, and Insecta (Supplementary Table S2). Of the 15,401,825 COI reads retained after quality filtering, 2,462,777 (16%) were classified by BOLDigger3 to the phylum Chordata, 2,797,463 (18%) to the phylum Arthropoda, with the remaining 10,141,585 (66%) of reads assigned to non-metazoan phyla such as fungi, bacteria, algae, or not assigned to any phyla.

Collectively with both markers, we detected 42 unique vertebrate taxa, 35 of which we assigned to a species known from Los Amigos, with the remaining seven taxa assigned to genus. Of the 105 beetles collected, we detected vertebrate DNA from 84 (80%) beetles (excluding human and domestic species). A mean of 2.26 vertebrate species was detected from each beetle (range 0-7). Primate DNA was detected from 53% of beetles, with rodent DNA detected from 26%, and carnivore DNA detected from 22%. Beetles from the genus *Sylvicanthon* yielded DNA from the greatest diversity of mammal species and the most arboreal species, followed by beetles from the genus Onthophagus (Table 4).

**Table 4.** Mammal detections by beetle genus from extraction-free iDNA metabarcoding of dung beetles from Los Amigos Biological Station, Madre de Dios, Peru.

|  |  | Yielded mammalian DNA |  | Number of species |  |  |  |
| --- | --- | --- | --- | --- | --- | --- | --- |
| Genus | Quantity | Number | Percent | All Mammal | Arboreal | Terrestrial | Volant |
| Sylvicanthon | 73 | 49 | 67% | 22 | 13 | 7 | 2 |
| Canthon | 10 | 9 | 90% | 7 | 4 | 3 | 0 |
| Onthophagus | 9 | 6 | 67% | 10 | 5 | 5 | 0 |
| Deltochilum | 5 | 3 | 60% | 3 | 2 | 1 | 0 |
| Dichotomius | 5 | 4 | 80% | 2 | 1 | 1 | 0 |
| Eurysternus | 3 | 3 | 100% | 6 | 3 | 3 | 0 |

Of the 105 beetle samples collected and sequenced, we retrieved Scarabaeidae COI sequences from 72 samples (69%). Taxonomic assignment by BOLDigger3 matched with morphological ID at the species level for 25 samples (35%) and at the genus level for 68 samples (94%; supplementary Table S3). The most common species we collected, *S. prosenii*, does not possess a publicly available reference sequence in BOLD, so it could only be identified at the genus level with BOLDigger3.

## Discussion

This study demonstrates that iDNA metabarcoding of dung beetles using an extraction-free, direct PCR approach is a powerful and efficient tool for surveying mammals in tropical forests. This approach was particularly successful in detecting primates and other arboreal species that can often be difficult to detect with traditional methods like camera trapping. By directly amplifying and sequencing mammal DNA left behind by beetles in tubes, we were able to bypass the laborious DNA extraction step used in most iDNA studies, greatly reducing processing time. This extraction-free approach has been used for carrion flies (Srivathsan et al. 2023, Rodgers et al. 2025), but to our knowledge it has never been applied to dung beetle metabarcoding. DNA extraction kits for environmental samples typically cost $3-$8 USD per sample, and thus even for small studies such as this one, our extraction free approach saves hundreds of dollars, and for larger studies, savings could quickly exceed $1000 USD. Additionally, dissection of individual beetle digestive tracts followed by typical DNA extraction workflows can take 3-6 hours per 24 samples, so extraction from hundreds or thousands of beetles can take days or even weeks.

From just 105 dung beetles, we detected 42 vertebrate species from 16 orders and 26 families. This included 33 mammal species and a remarkable 10 out of the 12 primate species known from Los Amigos, with 53% of beetle samples containing primate DNA. Among the primates detected were the near-threatened shock-headed capuchin *Cebus cuscinus* and the endangered Peruvian spider monkey *Ateles chamek*. We also detected the rare Rio Tapajós saki monkey *Pithecia irrorate*, which is seldom observed and is listed as data deficient by the IUCN (IUCN 2025). The endangered Peruvian spider monkey *Ateles chamek* was the most detected species (n=38 samples), supporting recent evidence that this species is abundant at Los Amigos (Pottie et al. 2025). The next most detected species were *South American coati Nasua nasua* (n=16), Toppin’s titi monkey *Plecturocebus toppini* (n=12) Red howler monkey *Alouatta seniculus* (N=12) and Hoffmann’s two-toed sloth *Choloepus hoffmanni* (n=10; Table 3). Black-headed night monkey *Aotus nigriceps*, a species which is difficult to visually survey due to its nocturnal nature, was also detected from nine samples.

In addition to primates, we detected numerous other arboreal taxa including rodents, a carnivore, a sloth, and five species of arboreal marsupials, three from the genus Marmosa and two from the genus Marmosops. These small marsupial genera are poorly studied because of their arboreal and nocturnal nature (Loretto and Vieira 2008). They are rarely detected by camera traps and are challenging to capture in live traps. This group is often less represented in reference sequence databases than other mammal groups, and we did not detect any of these species with our 16S marker. As of this writing, just one 16S sequence for the genus Marmosa is available on NCBI Genbank (none from the species we detected with COI) and none for the genus Marmosops. Because COI confirmed their presence, it is likely that DNA from these species were amplified by the 16S marker, but we were unable to classify the resulting sequences due to lack of references in available databases. With the COI marker, we were able to assign four of the five small marsupial taxa to species, with one OTU only classified to the genus Marmosops. It is most likely that this OTU belongs to a species of Marmosops that lacks a COI barcode, despite COI barcodes from five Marmosops species from Peru recently added to BOLD (Sánchez-Vendizú et al. 2025). However, as new species of Marmosops are still being described, and more species are likely to be discovered due to their cryptic behavior (Díaz-Nieto and Voss 2016, Ferreira et al. 2020) it is also possible that this OTU belongs to an undescribed species.

Although we detected an impressive number of vertebrate species, there were likely additional species that were missed due to lack of reference sequences. Reference sequences for hyper-diverse regions like Peru make up just a small fraction of databases compared to sequences from the global North (Sánchez-Vendizú et al. 2025). Thus, many vertebrate species, especially from small mammals and marsupials, lack reference sequences for Peru. Reference sequences are even lacking for some charismatic taxa including primates and carnivores. As mentioned earlier, reference sequences are not available for 16S or COI for the primate *Pithecia irrorate*, although we were able to infer detection of this species because we assigned an OTU to the genus *Pithecia, and Pithecia irrorate* is the only species from the genus *Pithecia* known from the region. There is also no reliable reference sequence for the short-eared dog *Atelocynus microtis;* A COI sequence is available on Genbank (Accession # AF028183); however, the sequence is extremely low quality, with many ambiguous bases, making it useless for taxonomic assignment. We detected an unknown canid OTU with both 16S and COI. Because there are reference sequences available from Bush dog *Speothos venaticus,* the other canid known from our study area, we believe the OTU we detected is likely from short-eared dog, but this could not be confirmed without a reference sequence. Thus, improving reference sequence representation for Peru should be a priority to improve future eDNA and iDNA monitoring in this extremely biodiverse region (Sánchez-Vendizú et al. 2025).

The work presented here builds on a foundation of research showing that vertebrate DNA can be obtained from dung beetle samples (Gillett et al. 2016, Gómez and Kolokotronis 2017). More recently, this method has been used as a survey method for mammals in tropical forests (Drinkwater et al. 2021), including in South America. Saranholi et al. (2024) detected 47 mammal OTUs from 403 dung beetles collected in the Brazilian Amazon through metabarcoding of dissected gut contents. Amongst the mammals they detected were six primate species, supporting our findings that dung beetle metabarcoding works well for primate surveys. Additionally, they found that mammal DNA was present in 70% of beetle samples, which is lower than the percentage of samples containing vertebrate DNA from our study (80%). This is notable, as it shows that our direct PCR extraction-free approach provided greater detection frequency than did gut dissection followed by DNA extraction. They also detected up to seven different mammal species from single beetles, the same maximum number per beetle that we detected.

We sequenced our samples using primers targeting two different mitochondrial markers, a marker designed specifically to amplify a fragment of the 16S rRNA gene from mammals and a more general marker designed to amplify the COI gene, the most commonly used gene for animal barcoding, from metazoans. These two markers are complimentary as they have contrasting strengths and weaknesses. Because the 16S marker is specific to mammals, it is very efficient and amplifies little bycatch from non-target species. We detected just one non-mammalian species with 16S, a fish species, although this marker is also known to occasionally amplify amphibian DNA (Rodgers et al. 2025). In our study, the 16S marker detected more mammal species than the COI marker (25 species for 16S vs. 22 species for COI) despite much lower sequencing depth. This specificity is also a weakness in other regards, however, because this marker is not expected to amplify DNA from non-mammalian vertebrate species that could be part of the dung beetle diet, or DNA from the dung beetles themselves. Another weakness of the 16S marker is that, although it is well represented in reference databases, it is not nearly as well represented as COI, especially for lesser studied groups like small mammals and marsupials.

The COI marker, conversely, is the best represented barcode marker for animals (Ratnasingham and Hebert 2007, Kress et al. 2015) which is a great strength. For example, as noted above, we did not detect any small arboreal marsupial species with the 16S marker, despite detecting them with COI. The COI marker we used is designed to amplify DNA from nearly all metazoans, which also allowed us to detect non-mammalian vertebrate species that were not targeted by the 16S marker, including the dung beetles themselves. However, this COI marker is not perfectly specific to metazoans as it also amplified considerable sequences from fungi and other non-target taxa. In fact, the majority of COI OTUs were from non-target species such as fungi and non-dung beetle arthropods. We anticipated this issue, and thus we sequenced our COI library with Illumina NextSeq on a run that produced nearly 50 times the number of sequences produced by the Illumina MiSeq run used for the 16S marker. Had we not sequenced our COI library at such depth, it is likely that we would have missed many vertebrate species, which highlights the importance of assessing assay specificity and expected biodiversity of samples when determining the appropriate sequencing depth for metabarcoding studies.

The high proportion of off-target, non-metazoan sequences observed in our COI dataset was potentially increased by primer modifications as well lab methodology. Of the COI reads retained after quality filtering, 16% were classified to the phylum Chordata, 18% to the phylum Arthropoda, and the remaining 66% of reads were assigned to non-target phyla such as fungi, algae, bacteria, or not assigned to any phyla by BOLDdigger3. For this study, we modified the already highly degenerate forward primer mlCOIintF (Leray et al. 2013) with an additional degeneracy at the 3’ end, specifically replacing a Y base (matches either C or T nucleotides) with an N base (matches all 4 nucleotides). This was done to better match all vertebrate species but may have also increased amplification of off-target taxa such as fungi. We note, however, that even the unmodified version of this primer is known for high rates of off-target amplification (Hajibabaei et al. 2019). We also ran PCR for the COI marker at the published annealing temperature of 46 °C (Leray et al. 2013) after observing no amplification at the manufacturer recommended temperature of 60 °C for direct PCR master mix, which may have further lowered specificity. Future studies may benefit from optimizing PCR annealing temperature to reduce off-target amplification with these modified COI primers.

For taxonomic assignment of the COI data, we examined two different taxonomic classifiers. We used the classifier also used on our 16S data, the Naïve Bayes classifier trained by scikit-learn (Pedregosa et al. 2011), as well as the classifier BOLDigger3 (Buchner and Leese 2020). BOLDigger3 is specifically designed for taxonomic assignment of COI data using the BOLD database (Ratnasingham and Hebert 2007), a database that is more rigorously curated (with every sequence tied to a voucher specimen) than other publicly available databases such as GenBank or MIDORI2, which is based on GenBank (Leray et al. 2022). BOLD does not possess reference sequences for the 16S gene, and thus it was not possible to use BOLDigger3 for the 16S data. For COI, results from both classifiers were highly similar for taxonomic classification of vertebrate species. BOLDigger3 did identify two mammal taxa not identified by the Naïve Bayes classifier, however, the Naïve Bayes classifier identified one bird species not identified by BOLDigger3. The Naïve Bayes classifier did assign a much greater number of OTUs at higher taxonomic levels (e.g. 89% of OTUs assigned to phylum by Naïve Bayes, vs, 11% for BOLDigger3). However, this is not unexpected, as BOLDigger3 only assigns OTUs if they have relatively high match percentage with a sequence from the BOLD database (in our case only OTUs that matched at >93%), whereas the Naïve Bayes classifier assigns OTUs even when match percentage is low. For dung beetle identification, BOLDigger3 performed far better than the Naïve Bayes classifier. We attribute this to the fact that our most collected dung beetle genus, *Sylvicanthon*, is not represented in GenBank or MIDORI2. Although BOLD does not possess publicly available reference sequences for our most collected species, *S. proseni*, it does possess sequences from our second most collected species, *S. bridarolli*, as well as sequences from several other species from the genus *Sylvicanthon*. By contrast, BOLDigger3 was able to assign OTUs from *Sylvicanthon* samples to the correct genus and sometimes species.

Use of the general metazoan COI marker allowed us to not only detect vertebrate species but also to amplify barcode sequences from the dung beetles themselves using our direct PCR approach. We were able to retrieve Scarabaeidae COI sequences from 69% of samples without the need for DNA extraction. In most cases, genetic ID matched morphological ID, at least at the genus level (Supplementary Table S2). Our most collected species based on morphological ID was *Sylvicanthon proseni*, which does not possess publicly available reference sequences in the BOLD or Genbank reference databases. Thus, nearly all sequences from samples identified morphologically as *S. proseni* were attributed by BOLDigger3 to the congener *S. aequinoctialis*, a species known from northern South America and Central America, but not Peru. Two samples identified by morphology as the species *Eurysternus caribaeus* were identified by COI as the congener *Eurysternus hypocrite.* As both species are known from Madre de Dios, it is possible that our morphological ID was incorrect. Samples identified by morphology as *Onthophagus osculatii* were assigned by BOLDigger3 to either *O. osculatii* or its congener *O. rubrescens*, which is known from northern South America but not Peru. In four cases, genetic ID did not match morphological ID at the genus level. In three of these cases, however, a beetle matching the genus from the genetic ID was captured in the same trap at the same time. Thus, it is possible that the mismatch in genetic vs. morphological ID was due to cross-contamination of DNA between beetles in the pitfall trap.

Despite these small discrepancies, we find it remarkable that we were able to obtain DNA sequences from both the beetles themselves, and DNA from their vertebrate dung meals, without the need for formal DNA extraction.

In seven cases, we captured dung-beetle species that were too large to fit into two mL microcentrifuge tubes. In these cases, we swabbed the beetles’ face and body with a cotton swab and then rubbed the cotton swab at the bottom of a two mL microcentrifuge tube to transfer DNA from the beetle to the tube. In four out of seven of these samples, we were able to amplify and sequence mammalian DNA, and in three of seven samples, we were able to amplify and sequence DNA from the beetles themselves. An advantage of this swabbing method is that beetles do not need to be kept in tubes for 24 hours, they can be swabbed and immediately released. Swabbing has been used successfully for prey metabarcoding from carrion beetles (Higdon et al. 2024), but not with extraction-free direct PCR. Future research should explore optimized methods for non-invasive, extraction-free metabarcoding of dung beetles using swabs.

COI metabarcoding detected a wide range of invertebrate species beyond dung beetles. We detected an astounding 1,634 non-dung beetle arthropod OTUs from 17 orders and 104 families. These results speak to the extremely high sensitivity of eDNA metabarcoding. Dung beetles appear to constantly encounter DNA from other invertebrate species. Additionally, our pitfall traps were filled with leaf litter, and many other invertebrate species found their way into traps, which may have accounted for some of the invertebrate DNA detected. Likewise, it is possible that some of the avian species we detected may not have been due to consumption of bird feces by beetles; it is possible that beetles encountered bird DNA from the leaf litter in the trap or from a bird perching and defecating on a trap. However, dung beetles are known to consume bird feces (Stavert et al. 2014) and Saranholi et al. (2024) detected numerous bird species from metabarcoding of dung beetle gut contents. Thus, we conclude that the bird DNA we detected likely resulted from beetles consuming bird feces and not from contamination.

We demonstrate here that small dung beetles such as those from the genera Sylvicanthon and Canthon are ideal candidates for mammal monitoring via iDNA, especially for arboreal species and primates. These small species are abundant in tropical forests, are easy to trap, and fit nicely into small tubes for simple and efficient extraction-free, direct-PCR metabarcoding. These species appear to readily gather DNA from arboreal mammal species, especially primates. We detected DNA from 10 primate species, as well as from arboreal carnivores, marsupials, rodents, and even bats. Thus, targeting small dung beetle species such as *Sylvicanthon* and *Canthon* for extraction-free metabarcoding is a highly efficient method for monitoring of mammals in tropical forests.

In addition to using dung beetle metabarcoding as a tool for surveying vertebrate diversity, our efficient methodology can also be applied to ecological research on dung beetle food webs (Gómez and Kolokotronis 2017, Pedersen et al. 2024). Understanding dung beetle food webs is important, as they provide essential ecosystem services including decomposition of animal feces, potentially reducing the spread of diseases and parasites that can negatively affect mammal and human populations (Nichols et al. 2008). For example, our study primarily captured beetles from the genus *Sylvicanthon*, a group of small dung beetles strongly associated with mature forests and dense canopy cover. Our results indicate that *Sylvicanthon* species are generalists, utilizing dung from a wide range of vertebrates including arboreal, terrestrial, and volant species (Table 4).

In conclusion, this study affirms the effectiveness of using dung beetle-derived iDNA for surveying mammals, and especially arboreal species, in tropical forests. Additionally, we present a key methodological innovation, an extraction-free, direct-PCR approach, which greatly streamlines and economizes lab work compared to more labor intensive past iDNA methods. By using two complimentary markers with direct PCR, we were able to survey a tropical vertebrate community while also obtaining sequences of the dung beetles themselves, offering a non-invasive and efficient approach for biodiversity monitoring.

## Acknowledgments

We thank the staff at Los Amigos Biological Station for making this work possible. We thank Sam Pottie for contributing expertise on Amazonian primate species. Funding for this work was provided by the Gordon and Betty Moore Foundation. This work was conducted on the ancestral lands of Indigenous peoples of the Madre de Dios region, including the Yine, Matsigenka, Harakbut, and others. We recognize their enduring cultural, spiritual, and ecological stewardship of these forests and the tremendous biodiversity they contain, and we honor their sovereignty and knowledge. Bionformatics analyses were performed using resources provided by the University of Utah Center for High Performance Computing.

## Notes

### Competing Interest Statement

The authors have declared no competing interest.

